# Tumor-selective effects of active RAS inhibition in pancreatic ductal adenocarcinoma

**DOI:** 10.1101/2023.12.03.569791

**Authors:** Urszula N. Wasko, Jingjing Jiang, Alvaro Curiel-Garcia, Yingyun Wang, Bianca Lee, Margo Orlen, Kristina Drizyte-Miller, Marie Menard, Julien Dilly, Stephen A. Sastra, Carmine F. Palermo, Tanner Dalton, Marie C. Hasselluhn, Amanda R. Decker-Farrell, Stephanie Chang, Lingyan Jiang, Xing Wei, Yu C. Yang, Ciara Helland, Haley Courtney, Yevgeniy Gindin, Ruiping Zhao, Samantha B. Kemp, Cynthia Clendenin, Rina Sor, Will Vostrejs, Amber A. Amparo, Priya S. Hibshman, Matthew G. Rees, Melissa M. Ronan, Jennifer A. Roth, Basil Bakir, Michael A. Badgley, John A. Chabot, Michael D. Kluger, Gulam A. Manji, Elsa Quintana, Zhengping Wang, Jacqueline A. M. Smith, Matthew Holderfield, David Wildes, Andrew J. Aguirre, Channing J. Der, Robert H. Vonderheide, Ben Z. Stanger, Mallika Singh, Kenneth P. Olive

## Abstract

Broad-spectrum RAS inhibition holds the potential to benefit roughly a quarter of human cancer patients whose tumors are driven by RAS mutations. However, the impact of inhibiting RAS functions in normal tissues is not known. RMC-7977 is a highly selective inhibitor of the active (GTP-bound) forms of KRAS, HRAS, and NRAS, with affinity for both mutant and wild type (WT) variants. As >90% of human pancreatic ductal adenocarcinoma (PDAC) cases are driven by activating mutations in *KRAS*, we assessed the therapeutic potential of RMC-7977 in a comprehensive range of PDAC models, including human and murine cell lines, human patient-derived organoids, human PDAC explants, subcutaneous and orthotopic cell-line or patient derived xenografts, syngeneic allografts, and genetically engineered mouse models. We observed broad and pronounced anti-tumor activity across these models following direct RAS inhibition at doses and concentrations that were well-tolerated *in vivo*. Pharmacological analyses revealed divergent responses to RMC-7977 in tumor versus normal tissues. Treated tumors exhibited waves of apoptosis along with sustained proliferative arrest whereas normal tissues underwent only transient decreases in proliferation, with no evidence of apoptosis. Together, these data establish a strong preclinical rationale for the use of broad-spectrum RAS inhibition in the setting of PDAC.

---

Activating mutations in the three RAS oncogene isoforms (HRAS, KRAS, and NRAS) are associated with approximately 20% of human cancers^1,2^. KRAS is the predominant isoform mutated in cancer, including in more than 90% of pancreatic ductal adenocarcinomas (PDAC), a leading cause of cancer mortality in the United States^3^ and globally. Though RAS proteins were long regarded as “undruggable” targets, recent advances have led to the development and approval of agents that target one specific RAS variant, KRAS^G12C 4,5^. This strategy has shown promising efficacy in *KRAS^G12C^*-mutant tumors, including the small fraction (∼1%) of PDAC cases harboring this allele^6,7^. However, resistance arises quickly in the majority of patients treated with KRAS^G12C^ inhibitors and various alterations that include secondary RAS mutations and receptor tyrosine kinase (RTK)-mediated activation of wild type RAS pathway signaling have been identified in patients who have progressed on these inhibitors^8–10^. The FDA-approved KRAS^G12C^ inhibitors, as well as two recently-described inhibitors (one targeting KRAS^G12D^ and one with broader KRAS specificity), selectively target the inactive, GDP-bound state of KRAS and are consequently vulnerable mechanisms of resistance that increase levels of GTP-bound KRAS including activation of upstream RTKs^11–13^. Here we evaluated the pharmacology and anti-tumor activity of a mechanistically distinct preclinical inhibitor, RMC-7977, with selectivity for the active, GTP-bound forms of all RAS isoforms, mutant and wild type, in multiple preclinical models of PDAC. Holderfield et al.^14^ describes the discovery of RMC-7977 (also known as RM-042) along with evidence that this agent can overcome some forms of acquired resistance to inhibitors that target RAS-GDP isoforms. A related RAS^MULTI^(ON) inhibitor, RMC-6236, is currently in early clinical evaluation (NCT05379985).

The singular role of mutant KRAS in PDAC oncogenesis has provoked myriad strategies for therapeutic intervention. Efforts to target prenylation of RAS proteins, upstream receptors, and downstream signaling were stymied by functional redundancies and compensatory feedback mechanisms^1,9,10,15,16^. Combinatorial strategies targeting multiple pathway effectors or compensatory responses can drive greater activity, but generally at the cost of reduced tolerability. Studies of *Kras^G12D^* gene deletion in engineered mouse models demonstrate that mutant KRAS plays an essential role in the maintenance of PDAC^17–19^. This was recently bolstered by preclinical evidence showing tumor regressions in PDAC models following pharmacologic inhibition of KRAS^G12D 11,12^. The development of RAS inhibitors with broader specificity has the potential to benefit the majority of PDAC patients while countering a wider range of resistance mechanisms. However, given the critical role of RAS proteins in normal tissue homeostasis^20,21^, a prevailing question is whether broad inhibition of RAS activity in tumors can be implemented with a suitable therapeutic index^22^.

## RMC-7977 exhibits broad anti-cancer activity in PDAC models

*KRAS* mutations in PDAC occur principally at codon 12 (*KRAS^G12X^*) with infrequent occurrence of mutations at codons 61 (6-7%) and 13 (1%)^23,24^. Consistent with the finding that cell lines with *KRAS^G12X^* mutations are particularly sensitive to RMC-7977^14^, we found that human PDAC cell lines were among the most sensitive in a large scale screen of 796 human tumor cell lines using the PRISM platform (Fig. 1a). In concentration-response cell viability assays, RMC-7977 exhibited low nanomolar potency in most human and murine PDAC cell lines, with reduced activity observed in a *KRAS^Q61H^* line and a *BRAF*-mutant line (Fig. 1b,c and Extended Data Fig. 1a). Similarly, RMC-7977 exhibited low nanomolar potency for growth inhibition of Matrigel-embedded human *KRAS^G12X^*-mutant PDAC patient-derived organoids (Fig. 1d).

**Fig 1.**
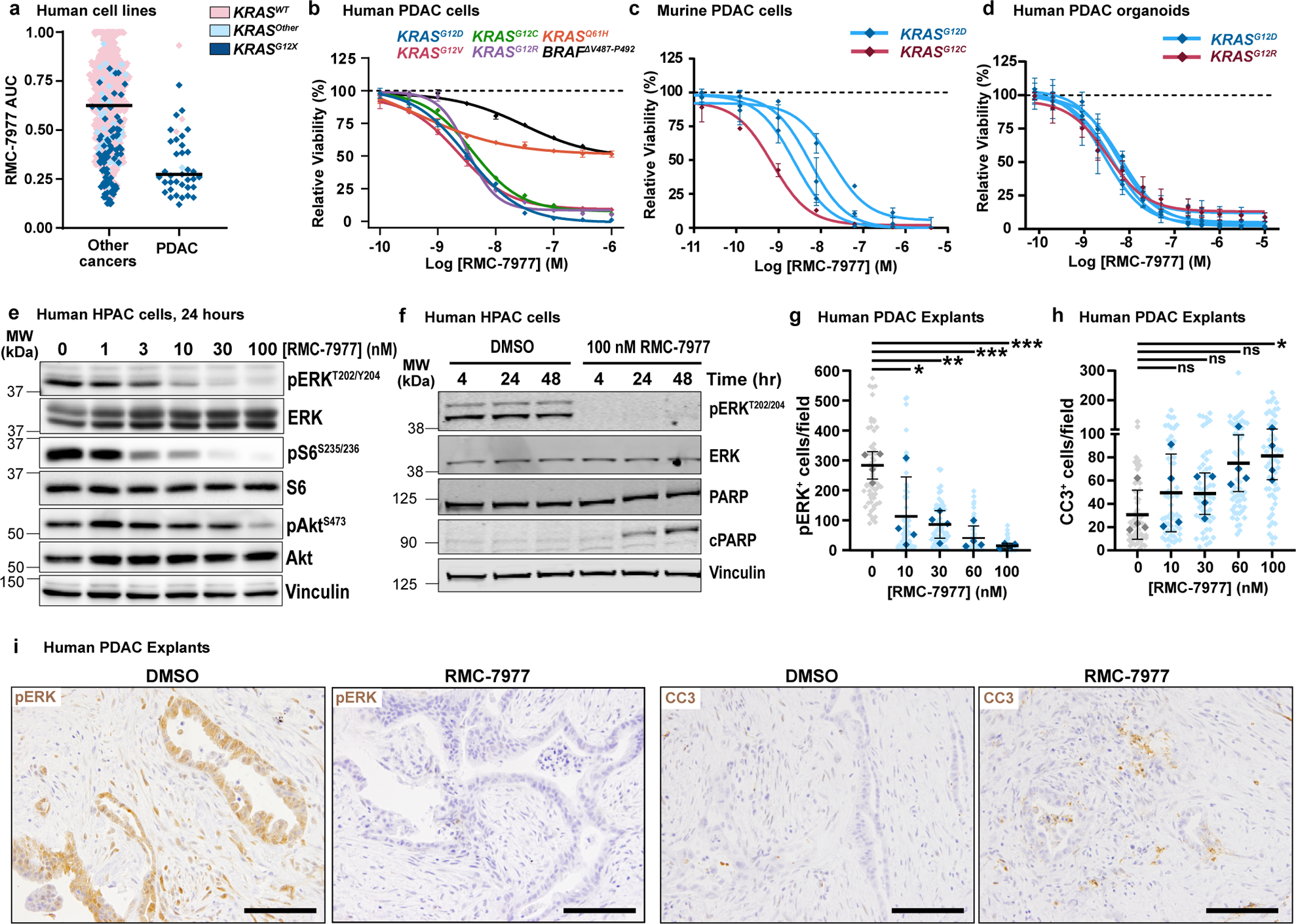
RMC-7977 demonstrates potent anti-tumor activity in *in vitro* and *ex vivo* models of PDAC. **(a)** PRISM multiplex cell line screening testing changes in viability of 796 cancer cell lines in response to RMC-7977 treatment. Cell line viability was plotted as Area Under Curve (AUC) values. KRAS status indicated by color of the symbol. Horizontal lines indicate median. **(b)** Viability levels of human PDAC cell lines with KRAS^G12X^ (HPAF-II, HuP-T4, MIA PaCa-2 and PSN-1), KRAS^Q61H^ (Hs 766T), and BRAF^ΔV487^–^P492^ (BxPC-3) mutations treated with indicated concentrations of RMC-7977 for 4 hours. **(c)** Viability levels of murine PDAC lines with KRAS^G12X^ (4662-G12D, 4662-G12C, 6419c5, 2838c3) mutations treated with indicated concentrations of RMC-7977 for 72 hours. **(d)** Viability levels of human PDAC organoids with KRAS^G12X^ mutations treated with indicated concentrations of RMC-7977 for 6 days. Data points in **(b, c, d)** represent the mean of technical 2-3 replicates normalized to DMSO control. Error bars indicate s.d. KRAS mutations are indicated by curve colors. **(e)** Western blots of HPAC cells treated with DMSO or range of RMC-7977 concentrations (1-100 nM) for 24 hours. Protein levels phospho-ERK^T202/204^, total ERK, phospho-pS6^S235/236^, total S6, phospho-Akt^S473^ and total Akt were analyzed. Vinculin was used as loading control. **(f)** Western blots of HPAC cells treated with DMSO or RMC-7977 (100 nM) for indicated time points. Protein levels phospho-ERK^T202/204^, total ERK, total PARP and cleaved PARP were analyzed. Vinculin was used as loading control. **(g-i)** *Ex vivo* human PDAC explants treated with DMSO or a range of RMC-7977 concentrations (10 – 100 nM) for 24 hr (n=4). **(g,h)** Quantification of IHC images of explants stained for **(g)** phospho-ERK^T202/204^ and **(h)** CC3. Analysis based on 5-15 fields of view (light shade), averaged per explant slice (dark shade) compared by one-way ANOVA test with Tukey correction (*, p<0.05; **, p<0.01; ***, p<0.001). Error bars indicate s.d. **(i)** Representative IHC images of phosho-ERK^T202/204^ and CC3 staining of explants treated with DMSO or RMC-7977 (100 nM). Scale bars = 50 μm.

Next, we evaluated RMC-7977 inhibition of RAS effector signaling. In human and murine PDAC cell lines, we observed reduced phosphorylation and activation of the RAS-RAF effector proteins ERK1/2 (pERK) at concentrations consistent with observed GI_50_ values (Fig. 1e and Extended Data Fig. 1b,c). In human cell lines, we detected more variable inhibition of PI3K effector signaling as monitored by phosphorylation of AKT (pAKT) and S6 (pS6), indicating some heterogeneity in the signaling responses across different lines (Fig. 1e and Extended Data Fig. 1c) and consistent with additional inputs (beyond direct RAS interactions) driving PI3K signaling in some contexts. Inhibition of pERK was generally durable over 48 hours in human cell lines and associated with induction of the apoptotic marker cleaved PARP (c-PARP) at later timepoints (Fig. 1f and Extended Data Fig. 1d).

Multiple studies have demonstrated that various stromal cells resident in PDAC tissues can modulate the sensitivity of malignant cells to therapy^25^. To assess the potency of RMC-7977 in the context of an intact, all-human microenvironment, we employed an *ex vivo* human PDAC explant model^26^, comprising cultured intact slices of freshly resected human PDAC tissue from patients at New York Presbyterian Hospital/Columbia University Irving Medical Center. Immunohistochemistry performed on PDAC explants treated for 24 hours with RMC-7977 showed a concentration-dependent decrease in pERK expression with an accompanying increase in the apoptosis marker, cleaved caspase 3 (CC3), with maximal changes at 100 nM, the highest concentration tested (Fig. 1g-i). These data are consistent with the pharmacodynamic responses observed in isolated cell line and 3D organoid systems and imply that the consequences of direct RAS inhibition observed *in vitro* are recapitulated in a complex PDAC tumor milieu. In summary, RMC-7977 consistently and potently inhibited RAS pathway signaling across *KRAS*-dependent PDAC cell lines, patient-derived organoids, and human tumor explants, resulting in growth attenuation and/or induction of apoptosis.

We next evaluated the activity of RMC-7977 *in vivo*, in a panel of seven different human PDAC cell line-derived xenografts (CDXs), as well as three fragment-based patient-derived xenograft (PDX) models of human PDAC, implanted either subcutaneously (SC) or orthotopically in the pancreas (Ortho) of immune-deficient mice (see Extended Data Table 1 for information on tumor models). RMC-7977 was administered at a daily dose of 10 mg/kg over the course of 21-28 days and resulted in significant anti-tumor activity in all 10 models, with tumor regressions of 30-98% relative to baseline in 7/10 models (Fig. 2a-d, Extended Data Fig. 2a-i). Critically, RMC-7977 was well tolerated in all 10 models, with treated animals generally exhibiting stable body weight over time (Fig. 2e, Extended Data Fig. 2j-r).

**Fig 2.**
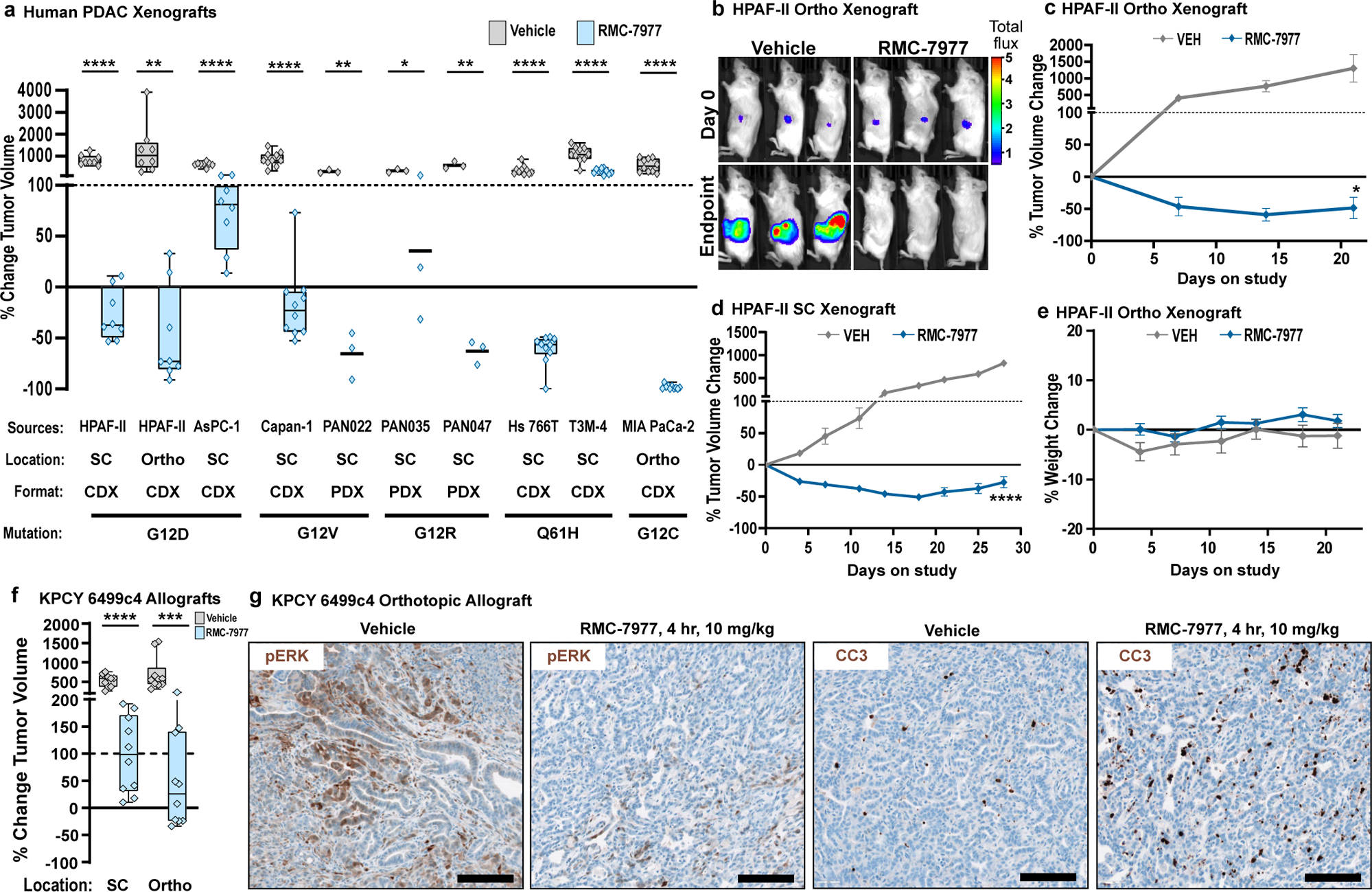
RMC-7977 exhibits anti-tumor activity in xenograft and allograft models of PDAC. **(a-e)** Human PDAC xenograft models implanted either subcutaneously (SC) or orthotopically (Ortho) into immunodeficient mice. Tumor-bearing mice were treated with Vehicle or RMC-7977 (10 mg/kg, po, q.d.; n=3-10) for 21-28 days. **(a)** Boxplot showing percent change in tumor volumes at endpoint compared with baseline at Day 0 in Vehicle and RMC-7977 treatment arms. Each symbol represents one mouse. Source and format of each cell line, KRAS mutation, and tumor locations are indicated in the graph. Study arms were compared by Student’s unpaired t-test (*, p<0.05; **, p<0.01; ****, p<0.0001). **(b)** Representative bioluminescence images showing signal in HPAF-II orthotropic xenograft tumors at Day 0 and at Day 21 for Vehicle and RMC-7977 treatment arms. **(c,d)** Representative tumor growth curves for HPAF-II **(c)** orthotopic and **(d)** subcutaneous xenograft models treated with Vehicle or RMC-7977 (n=8/group), shown as percent tumor volume change from baseline over the course of treatment. Vehicle and RMC-7977 groups were compared by 2-way repeated measures ANOVA on the last measurement day of the vehicle group (*, p<0.05; ****p<0.0001). Error bars indicate ± s.e.m. **(e)** Tolerability of RMC-7977 as assessed by percent animal body weight change from baseline over the course of treatment. Error bars indicate ± s.e.m. **(f, g)** KPCY-derived PDAC cell line (6499c4) was transplanted either subcutaneously or orthotopically into syngeneic mice. Tumor-bearing mice were treated with Vehicle or RMC-7977 (10 mg/kg, po, q.d.; n=9-10/group). **(f)** Boxplot showing changes in subcutaneous and orthotopic tumor volumes at Day 14, compared with baseline at Day 0, in Vehicle and RMC-7977 treatment arms. Groups compared by Student’s unpaired t-test (***, p<0.001; ****, p<0.0001). Tumor locations as indicated in the graph. **(g)** Representative IHC images of Vehicle and RMC-7977-treated KPCY allograft tumors stained for phospho-ERK^T202/204^ and CC3. Scale bars = 100 μm.

Next, we examined the anti-tumor activity of RMC-7977 in C57Bl/6 mice with intact immune systems by evaluating cell-line derived allografts (CDAs) implanted either subcutaneously or orthotopically with a PDAC cell line from *the Kras^LSL.G12D/+^*; *p53^R172H/+^*; *Pdx1-Cre^tg/+^*; *Rosa26^LSL.YFP/+^* (KPCY) mouse model on a matched background. In this setting, RMC-7977 diminished tumor growth and extended overall survival, though without inducing significant regressions (Fig. 2f and Extended Data Fig. 2s,t). Pharmacodynamic analysis of the orthotopic CDA model showed reduced expression of pERK and increased apoptosis in the tumors four hours after treatment, followed by recovery of pathway activity at 24 hours (Fig. 2g and Extended Data Fig. 2u). Quantitative analysis of RMC-7977 levels in matched tumor tissue samples found that the restoration of RAS/MAPK activity in these tissues was consistent with the observed tumor half-life of 3.5 hours in this model (Extended Data Table 2).

## The pharmacology of RAS addiction

To understand the mechanistic basis of the broad anti-tumor effects described above, we carried out a detailed and quantitative analysis of the pharmacological profile of RMC-7977 in the setting of PDAC. We first examined the association of drug concentration and RAS pathway inhibition in a representative human CDX model of PDAC (Capan-1; *KRAS^G12V^*). We assessed the response of tumor cells to treatment with a single dose of 10, 25, or 50 mg/kg RMC-7977 by measuring expression of human *DUSP6*, a RAS/MAPK pathway transcriptional target, in tumor tissues by qRT-PCR. *DUSP6* levels were effectively inhibited for 24-48 hours post-dose and pathway inhibition was tightly associated with concentrations of RMC-7977 in tumors (Fig. 3a,b; EC_50_ = 142 nM in Capan-1 CDX tumors), indicating that the local total drug concentration is a critical determinant of biochemical activity. We also observed somewhat prolonged RMC-7977 exposure in Capan-1 xenograft tumors, with a ∼3-fold increase in overall exposure (AUC_0-48_) in tumors compared to that in blood (Extended Data Table 2).

**Fig 3.**
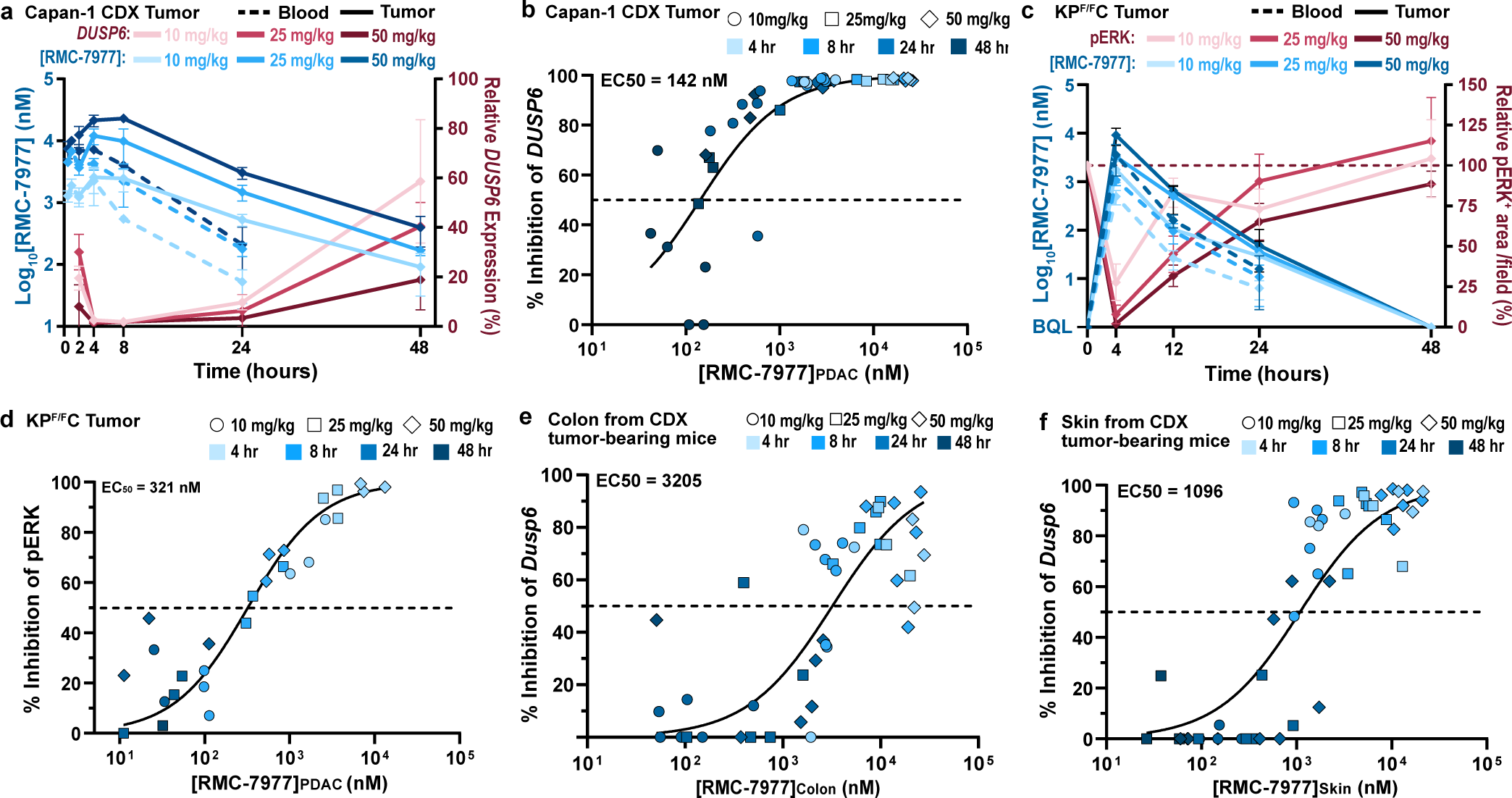
Pharmacology of RAS addiction. **(a,b)** Capan-1 subcutaneous xenograft tumors were treated with Vehicle or RMC-7977 at 10 mg/kg, 25 mg/kg and 50 mg/kg. Tissues were harvested at indicated timepoints (n=3-6/timepoint/dose). **(a)** PK/PD in the Capan-1 xenograft model. Pharmacokinetic profile shown as RMC-7977 concentration in tumors (solid blue lines) and blood (dashed blue lines) over time. Pharmacodynamic response shown as relative change in *DUSP6* mRNA expression (solid red lines). Shades of blue or red represent three tested doses. Values plotted as mean ± s.d. **(b)** PKPD relationship between RMC-7977 concentration and inhibition of *DUSP6* expression in tumors. **(c,d)** Tumor-bearing KP^F/F^C mice were treated with RMC-7977 at 10 mg/kg, 25 mg/kg and 50 mg/kg. Tissues were harvested at indicated timepoints (*n*=3/timepoint/dose). **(c)** PK/PD in the tumors isolated from KP^F/F^C mice. Pharmacokinetic response shown as RMC-7977 concentration in tumors (solid blue lines) and blood (dashed blue lines) over time. Pharmacodynamic response shown as relative change in pERK positive IHC staining in tumors (solid red lines) over time. Shades of blue or red represent three tested doses. Values plotted as mean ± s.d. **(d)** PKPD relationship between RMC-7977 concentration and pERK inhibition in tumors. **(e,f)** PKPD relationship between RMC-7977 concentration and inhibition of *Dusp6* expression in normal **(e)** colons and **(f)** skin, isolated from CDX tumor-bearing mice. **(b,d,e,f)** A 3-parameter sigmoidal exposure response model was fitted to the data to derive EC_50_ values. Timepoints represented by shades of blue and doses represented by symbol shapes.

The delivery of drugs to autochthonous PDAC tissues can be impeded by its expansive desmoplastic stroma, which both suppresses tumor vascularity and impedes diffusion^27,28^. To assess the impact of a native stromal and immune microenvironment on the pharmacology of RMC-7977, we utilized *Kras^LSL.G12D/+^*; *p53^Flox/Flox^*; *Pdx1-Cre^tg/+^* (KP^F/F^C) mice^29^, a genetically engineered model that rapidly develops autochthonous *Kras^G12D^* mutant PDAC. We observed that pathway modulation in the KP^F/F^C model was less sensitive to RMC-7977 as compared to the Capan-1 CDX model and required higher doses (25 to 50 mg/kg) to fully suppress RAS/MAPK signaling (Fig. 3a,c and Extended Data Fig. 3a). Examining the pharmacokinetic profile of RMC-7977 in KP^F/F^C mice, we observed relatively lower exposure in blood and PDAC tumors compared to CDX tumor-bearing BALB/c immune-deficient mice, comparable to that in the KPCY CDA model, across multiple dose levels. This implies that strain- and model-specific variables could contribute to the exposure and relative activity of RMC-7977 (Fig. 3a,c and Extended Data Table 2). We also observed a shorter half-life (t_1/2_) of RMC-7977 in KP^F/F^C tumors compared to CDX tumors and concordantly faster recovery of pERK levels. Importantly, the tight relationship between drug concentration and pathway suppression observed in CDX models was maintained in this autochthonous model (Fig. 3d, EC_50_ = 321 nM). Moreover, in all model systems tested, concentrations of RMC-7977 were higher in tumor tissues than in blood (K_p_ > 1, Extended Data Table 2). Based on this and the reproducible tumor concentration/response (PK/PD) relationship across models, we predicted that daily or alternate day dosing schedules would yield an effective and metronomic pattern of RAS/MAPK pathway suppression in pancreatic tumors.

The broad anti-cancer activity of RMC-7977 in a wide range of preclinical models of PDAC at tolerable dose levels raises the question of how normal tissues survive the inhibition of RAS signaling. To explore this, we first examined the concentration-response relationship of RMC-7977 for murine *Dusp6* mRNA inhibition (via qRT-PCR) in the normal colon and the skin, two proliferative tissue compartments known to rely on RAS signaling for self-renewal^30,31^. RMC-7977 demonstrated appreciably greater potency for RAS pathway modulation in tumor cells of the Capan-1 CDX model (EC_50_ = 142 nM, Fig. 3b) than in these normal tissues, exhibiting a ∼22 fold and ∼8 fold lower EC_50_ compared to colon and skin, respectively (Fig. 3b,e,f and Extended Data Fig. 3b,c). We also examined pERK suppression via RMC-7977 in colon tissues from the KP^F/F^C cohort and found that RAS/MAPK activity was rapidly restored to baseline at the 10 and 25 mg/kg dose levels while pathway suppression was more prolonged in tumors at these doses (Fig. 3c and Extended Data Fig. 3d,e). Together these data highlight a marked difference in the potency and kinetics of RMC-7977-mediated pharmacodynamic pathway modulation in normal tissues expressing RAS^WT^ compared to KRAS^G12X^ driven PDAC tumors^30^.

We then examined the downstream consequences of RAS inhibition in both tumor and normal tissues. In Capan-1 CDX tumors, we observed a notable increase in CC3^+^ apoptotic cells, peaking at 8 hours post-dose, relative to vehicle controls. By contrast, few apoptotic cells were observed in the colon or the skin at any timepoint (Fig. 4a,b). In KP^F/F^C tumors, we observed a sharp, ∼4-fold spike in CC3^+^ apoptotic cells at four hours following a single dose of RMC-7977, that was not observed in the matched colon tissues from these mice (Fig. 4c,d). In both the KP^F/F^C and CDX models, the kinetics of apoptosis initiation in tumors closely mirrored the full inhibition of pERK. In stark contrast, apoptosis in normal tissues was negligible or absent across all doses and time points, even at the times when pERK was fully inhibited.

**Fig 4.**
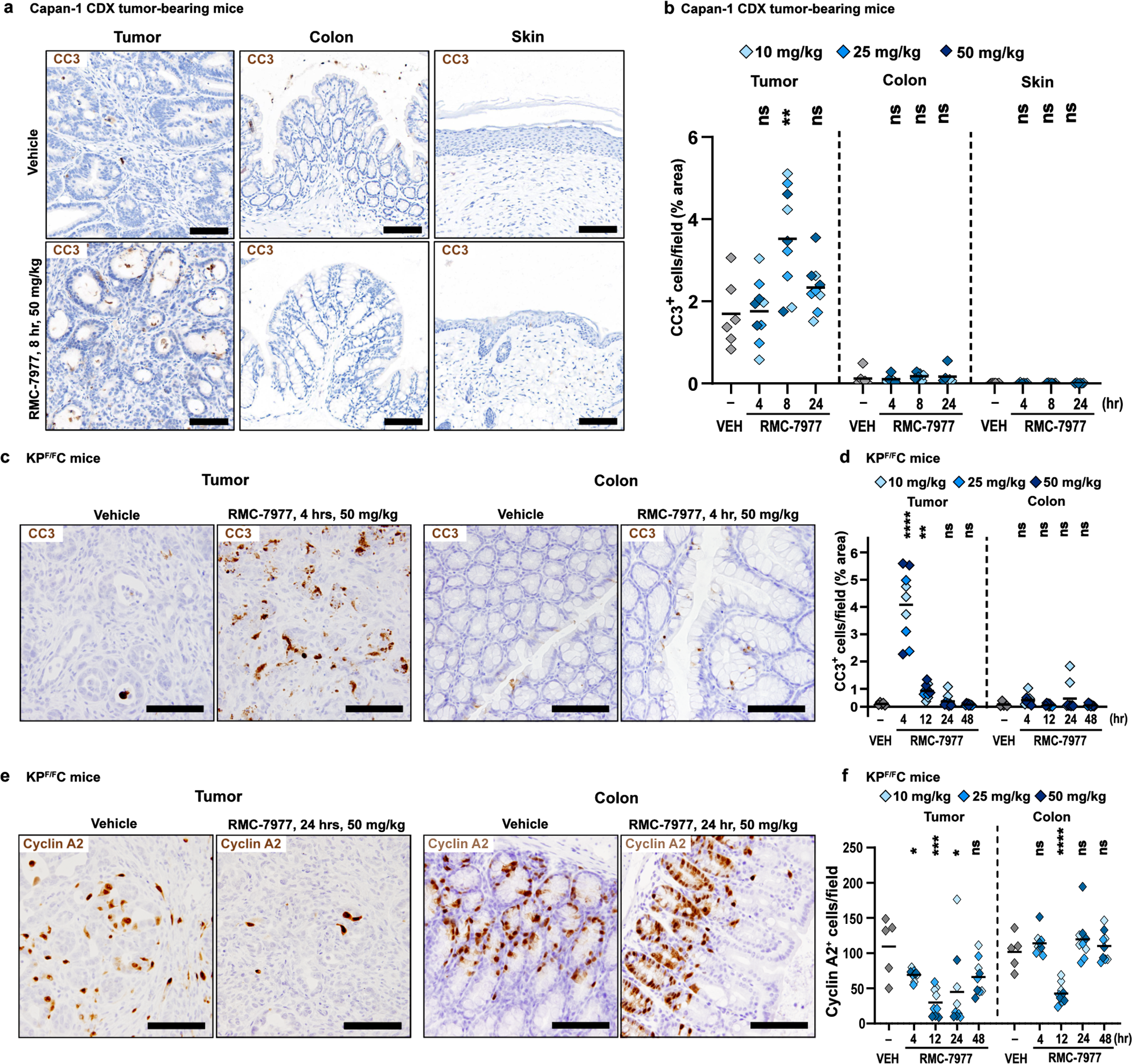
Inhibition of RAS induces pancreatic tumor-selective apoptosis. **(a)** Representative IHC images of tumors, colons, and skin from Capan-1 xenograft model collected at 8 hours post single dose of Vehicle or RMC-7977, stained for CC3. Scale bars = 100 μm. **(b)** Quantification of CC3 staining in tumors, colon, and skin. **(c)** Representative IHC images of KP^F/F^C tumors and colons collected at 4 hours post single dose of Vehicle or RMC-7977, stained for CC3. Scale bars = 50 μm. **(d)** Quantification of CC3 staining in tumors and colons. **(e)** Representative IHC images of KP^F/F^C tumors and colons collected at 24 hours post single dose of Vehicle or RMC-7977, stained for Cyclin A2. Scale bars = 50 μm. **(f)** Quantification of Cyclin A2 staining in tumors and colons. **(b,d,f)** Analysis of IHC images based on 10-15 fields of view and plotted as average per tissue section. Shades of blue represent three tested doses. Results were compared by Student’s unpaired t-test (*, p<0.05; **, p<0.01; ***, p<0.001; ****, p<0.0001).

Finally, we detected a sustained decrease in cell proliferation in KP^F/F^C tumors (as measured by IHC for Cyclin A2) beginning 4 hours after treatment with RMC-7977, with maximal inhibition maintained at 12 and 24 hours and a partial recovery at 48 hours (Fig. 4e,f). In contrast, in matched colon tissue from these KP^F/F^C mice, we observed only a transient decrease in proliferation at 12 hours, with Cyclin A2 levels fully restored at 24 hours (Fig. 4e,f). Thus, the overall proliferation of this self-renewing normal tissue was minimally impacted compared to the sustained anti-proliferative response observed in PDAC tissues. Together, these results demonstrate key differences in how RAS^WT^ normal tissues react and adapt to metronomic RAS inhibition with a broad-spectrum RAS inhibitor compared to tumors driven by mutant KRAS, providing a rational basis for the tumor-selectivity of RAS inhibition in PDAC.

## Preclinical efficacy of RMC-7977 in autochthonous models of PDAC

To evaluate the anti-tumor activity of RMC-7977 in clinically predictive models of human PDAC, we first carried out short-term intervention studies in two variants of the *Kras^LSL.G12D/+^; p53^LSL.R172H/+^; Pdx1-Cre^tg/+^* (KPC) model. Performed independently at separate institutions, the first study treated a YFP-lineage traced version of the KPC model (KPCY), on a pure C57Bl/6 background, with daily 25 mg/kg RMC-7977, for two weeks (Fig. 5a-d). The second study treated KPC mice, on a background enriched for 129S_4_/SvJae, with 50 mg/kg RMC-7977 q.o.d., for one week (Fig. 5e-k). In both experiments, RMC-7977 treatment induced stable disease or regressions in most tumors with little impact on animal weight (Fig. 5b-d,f-h).

**Fig 5.**
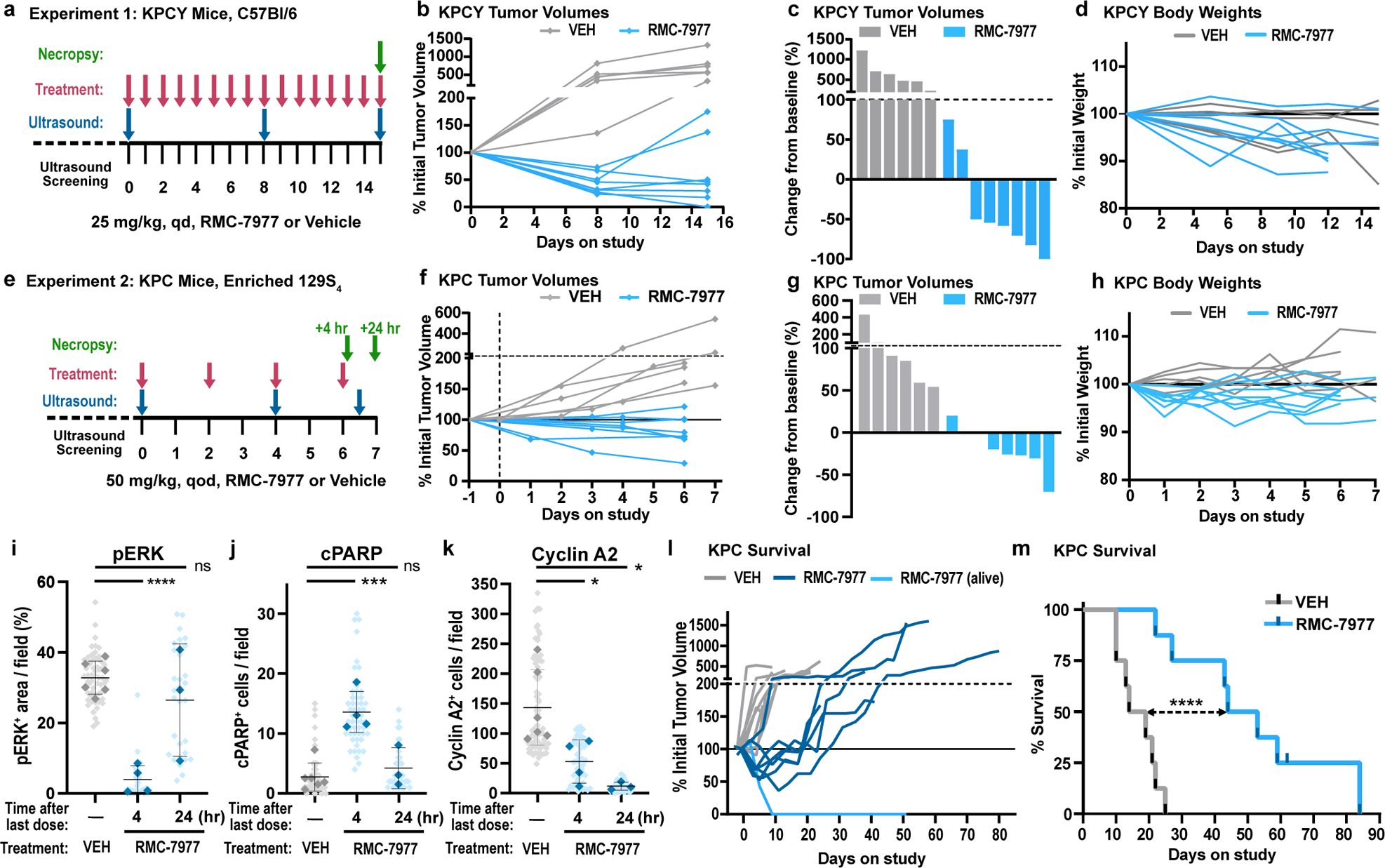
RMC-7977 inhibits tumor growth and extends survival in autochthonous models of PDAC. **(a)** Schematic of KPCY mice (on C57Bl/6 background) treatment with Vehicle (n=6) or RMC-7977 (25 mg/kg, po, q.d.; n=8) for 15 days. **(b)** Tumor growth curves for mice in experiment as depicted in panel (a). Each line represents one mouse and each symbol represents ultrasound scan. **(c)** Waterfall plot showing percent change in tumor volume compared to baseline of KPCY after 15 days of treatment. **(d)** RMC-7977 tolerability as assessed by animal body weight change over the course of treatment. **(e)** Schematic of KPC mice (on 129S_4_/SvJae background) treatment with Vehicle (n=6) or RMC-7977 (50 mg/kg, po, q.o.d.; n=8) for 1 week. Tissues collected at 4 hours (n=4) or 24 hours (n=4) post last dose. **(f)** Tumor growth curves for mice in experiment as depicted in panel (e). Each line represents one mouse and each symbol represents ultrasound scan. **(g)** Waterfall plot showing percent change in tumor volume compared to baseline after 1 week of treatment. **(h)** RMC-7977 tolerability assessed by animal body weight change from baseline over the course of treatment. **(i-k)** IHC analysis of KPC tumors. Tumors stained for **(i)** phospho-ERK^T202/204^; **(j)** cleaved PARP, **(k)** cyclin A2. Quantification of IHC images was compared between Vehicle, RMC-7977 (1 week + 4 hour) and RMC-7977 (1 week + 24 hour). Quantification was based on 10-15 fields of view (light shade), averaged per tumor (dark shade) and means were compared by Student’s unpaired t-test (*, p<0.05; ***, p<0.001; ****, p<0.0001). Error bars indicate s.d. (**l,m**) KPC mice treated with Vehicle (n=8) or RMC-7977 (50 mg/kg, po, q.o.d., n=8) until endpoint criteria were met. (**l**) Tumor growth rates calculated from longitudinal tumor volumes. (**m**) Kaplan-Meier survival analysis comparing RMC-7977 and Vehicle (historical) treatment arms (****, p<0.0001).

Immunohistochemical analyses of tumor tissues from KPC mice from the second study revealed decreased pERK expression and increased expression of the apoptotic markers c-PARP and CC3 in tumors collected four hours after the final dose (Fig. 5i,j and Extended Data Fig. 4a). pERK, c-PARP, and CC3 returned to baseline in tumors collected 24 hours after the last dose, consistent with results from the single dose study in KP^F/F^C mice (Fig. 4d). These data suggest that the metronomic effect of RMC-7977 on pERK and apoptosis in tumors persists after a week of treatment and that each successive dose induces an additional wave of apoptosis. Additionally, we observed a persistent decrease in proliferation (by Cyclin A2 IHC) in RMC-7977 treated KPC tumors, with maximal inhibition at 24 hours after the final dose (Fig. 5k), consistent with results from the single dose study in KP^F/F^C mice above (Extended Data Fig. 3f). Together, these results indicate that RAS inhibition has a cyclical impact on the induction of apoptosis in pancreatic tumors and a sustained effect on tumor proliferation during the initial week of treatment.

Next, we performed an interventional survival study in tumor-bearing KPC mice. Remarkably, animals treated with 50 mg/kg RMC-7977, q.o.d. exhibited tumor stabilizations and regressions as measured by longitudinal, high-resolution 3D-ultrasound^32^. By contrast, tumors in vehicle-treated mice uniformly exhibited progressive growth (Fig. 5l). Kaplan-Meier analysis of overall survival showed a striking ∼3-fold increase in median survival in the RMC-7977 treated cohort as compared to controls (Fig. 5m), far exceeding the most effective monotherapy tested in the KPC model^33^. The body weights of KPC mice treated with RMC-7977 were similar to those of control KPC mice (Extended Data Fig. 4b). Together, these data demonstrate the preclinical anti-tumor activity and tolerability of RMC-7977 in clinically relevant mouse models and highlight the potential of broad-spectrum RAS inhibition to offer clinical benefit to patients with *KRAS*-mutant PDAC.

The alluring possibilities of a mutation-agnostic RAS inhibitor as a therapeutic agent for RAS-addicted cancers have been inseparably entangled with the widely held assumption that targeting wild-type RAS would prove intolerable. Indeed, there was previouly little direct evidence available from mouse models to guide expectations for the effects of widespread inhibition of canonical RAS family members in mature tissues. Homozygous knockout of *Kras* produces an early (e3.5) embryonic lethal phenotype^20^ and conditional deletion in hematopoietic lineages eventually compromises hematopoiesis^21^. However, neither of these experiments accurately model the acute inhibition of RAS in adults that can be achieved with a broad-spectrum RAS inhibitor. Perhaps the closest parallel is the systemic inhibition of C-MYC (which serves as a conduit for RAS signaling in many cell types) through the inducible expression of the dominant negative protein Omomyc^34^. This approach showed that inhibition of physiological MYC activity reduced proliferation in most epithelial tissues, but that key epithelial functions were broadly maintained for extended periods of time. Systemic RAS inhibition may prove to be similar in nature, with tolerability enabled by the relatively low levels of active RAS-GTP (the target for RMC-7977) in normal tissues^35^ and the somewhat reduced affinity of RMC-7977 for wild type RAS as compared to mutant variants^14^. This is consistent with prior work showing that normal cells can rapidly restore homeostasis following RAS pathway inhibition in contrast to RAS-addicted tumor cells^36^ and that normal RAS^WT^ skin cells behave differently following wounding stimuli as compared to RAS^MUT^ skin cells^37^. By contrast, the anti-proliferative, pro-apoptotic effects of RMC-7977 in KRAS^MUT^ tumor cells and tissues are consistent with the concept of “oncogene addiction” and explain the remarkable extension of overall survival we observed in the highly chemoresistant KPC mouse model. Taken together, our findings have significant and positive implications for the translation of broad-spectrum RAS inhibition as a monotherapy in patients with pancreatic ductal adenocarcinoma and potentially other types of RAS-addicted cancers.

## Material and Methods

### RMC-7977 formulation

For *in vitro* studies RMC-7977 was re-suspended in DMSO (Fisher Bioreagents, BP231-100) and used at 10 mM stock concentration. For use in the *in vivo* studies RMC-7977 was prepared using the formulation made of 10/20/10/60 (%v/v/v/v) DMSO/PEG 400/Solutol HS15/water. The same vehicle formulation was used for all control groups.

### Cell culture and reagents

All human cancer cell lines were obtained from American Type Culture Collection (ATCC) and grown in appropriate medium supplemented with 10% fetal bovine serum (FBS) and 1% penicillin/streptomycin and maintained at 37°C in a humidified incubator at 5% CO_2_, unless otherwise indicated. PDAC cell lines generated from patient-derived xenografts (Pa01C, Pa14C, and Pa16C) were a gift of Dr. Anirban Maitra (MD Anderson Cancer Center).

Murine PDAC cell lines were derived from tumor-bearing KPCY (6419c5, 2838c3^38^) or KPC (4662-G12D^39^) animals on a congenic C57BL/6 background. The 4662-G12C line was generated using CRISPR/Cas9 to replace the endogenous G12D mutation from 4662-G12D cells with the G12C mutation by lentiviral transduction. KRAS allele states were confirmed by genomic sequencing. Cells were cultured in Dulbecco’s modified Eagle medium (DMEM, high glucose without sodium pyruvate) supplemented with 10% heat-inactivated FBS and 1% penicillin/streptomycin. (See Supplementary Methods Table 1 for cell line information)

### PRISM assay

#### Cell Lines

The PRISM cell set consisted of 796 cell lines representing more than 45 lineages (See Supplementary Methods Table 2 for cell line information), which largely overlapped with the Cancer Cell Line Encyclopedia (CCLE) (https://portals.broadinstitute.org/ccle). Cell lines were grown in RPMI without phenol red, supplemented with 10% or 20% FBS for adherent and suspended lines, respectively. Parental cell lines were stably infected with a unique 24-nucleotide DNA barcode via lentiviral transduction and blasticidin selection. After selection, barcoded cell lines were expanded and subjected to quality control (mycoplasma contamination test, a SNP test for confirming cell line identity, and barcode ID confirmation). Approved cell lines were then pooled (20-25 cell lines per pool) based on doubling time similarity and frozen in assay-ready vials.

#### PRISM Screening

RMC-7977 was added to 384-well plates at 8-point concentration with 3-fold dilutions in triplicate. These assay-ready plates were then seeded with the thawed cell line pools. Adherent cell pools were plated at 1250 cells per well, while suspension and mixed adherent/suspension pools were plated at 2000 cells per well. Treated cells were incubated for 5 days, then lysed. Lysate plates were collapsed together prior to barcode amplification and detection.

#### Barcode Amplification and Detection

Each cell line’s unique barcode is located in the 3’UTR of the blasticidin resistance gene and therefore is expressed as mRNA. Total mRNA was captured using magnetic particles that recognize polyA sequences. Captured mRNA was reverse-transcribed into cDNA and then the sequence containing the unique PRISM barcode was amplified using PCR. Finally, Luminex beads that recognize the specific barcode sequences in the cell set were hybridized to the PCR products and detected using a Luminex scanner which reports signal as a median fluorescent intensity (MFI).

#### Data Processing

I. Each detection well contained 10 control barcodes in increasing abundances as spike-in controls. For each plate, we first create a reference profile by calculating the median of the log_2_(MFI) values across negative control wells for each of these spiked-in barcodes.
II. For each well, a monotonic smooth p-spline was fit to map the spike in control levels to the reference profile. Next, we transform the log_2_(MFI) for each cell barcode using the fitted spline to allow well-to-well comparisons by correcting for amplification and detection artifacts.
III. Next, the separability between negative and positive control treatments was assessed. In particular, we calculated the error rate of the optimum simple threshold classifier between the control samples for each cell line and plate combination. Error rate is a measure of overlap of the two control sets and was defined as Error = (FP + FN)/n, where FP is false positives, FN is false negatives, and n is the total number of controls. A threshold was set between the distributions of positive and negative control log_2_(MFI) values (with everything below the threshold said to be positive and above said to be negative) such that this value is minimized. Additionally, we also calculated the dynamic range of each cell line. Dynamic range was defined as DR=μ_-_ - μ_+_, where μ+/− stood for the median of the normalized logMFI values in positive/negative control samples.
IV. We filtered out cell lines with error rate above 0.05 or a dynamic range less than 1.74 from the downstream analysis. Additionally, any cell line that had less than 2 passing replicates was also omitted for the sake of reproducibility. Finally, we computed viability by normalizing with respect to the median negative control for each plate. Log-fold-change viabilities were computed as log-viability = log_2_(x) – log_2_(μ−), where log_2_(x) is the corrected log_2_(MFI) value in the treatment and log_2_(μ−) is the median corrected log_2_(MFI) in the negative control wells in the same plate.
V. Log-viability scores were corrected for batch effects coming from pools and culture conditions using the ComBat algorithm^40^.
VI. We fit a robust four-parameter logistic curve to the response of each cell line to the compound: f(x) = b + (a – b)/(1 + e^s log(x/EC50)^) With the following restrictions:

1. We require that the upper asymptote of the curve be between 0.99 and 1.01
2. We require that the lower asymptote of the curve be between 0 and 1.01
3. We do not enforce decreasing curves
4. We initialize the curve fitting algorithm to guess an upper asymptote of 1 and a lower asymptote of 0.5
5. When the standard curve fit fails, we report the robust fits provided by the dr4pl R-package

and computed AUC values for each dose-response curve and IC_50_ values for curves that dropped below 50% viability.
VII. Finally, the replicates were collapsed to a treatment-level profile by computing the median log-viability score for each cell line.

#### Associations between inhibitor sensitivity Area Under the Curve (AUC) and mutations

For every gene with non-silent mutations in at least four cell-lines, we compared the AUC values between cells with and without those mutations using a t-test. This analysis was carried out for: (i) the full dataset; (ii) excluding cell lines with non-silent KRAS mutations; and (iii) excluding cell lines that have either KRAS or NRAS non-silent mutations.

#### Bioinformatics analyses

Gene mutation, gene expression, and lineage data were downloaded from the 22Q4 release of the DepMap Data Portal^41^. For tumor models with no publicly available data, we carried out whole exome sequencing to ascertain gene mutations and RNA sequencing to ascertain gene expression. DNA mutation calling was accomplished with TNSeq using the hg38 version of the human genome^42^. Functional annotation of the resulting mutation calls was accomplished with Variant Effect Predictor and further annotated with oncoKB^43^. Gene expression was quantified using salmon against the hg38 version of human transcriptome further processed using txImport and edgeR to generate normalized counts^44–46^.

### Murine cell viability assays

PDAC murine cell lines harboring *Kras^G12D^* or *Kras^G12C^* mutations were seeded at 2 × 10^3^ in a 96-well plate. Cells were treated 24 hours later with DMSO or serial dilutions of RMC-7977. Cell viability was evaluated 72 hours later by measuring adenosine triphosphate (ATP) levels using the CellTiter-Glo® Luminescent Cell Viability Assay (Promega, G7572) according to the manufacturer’s instructions. Technical triplicates were run for each biological replicate and a total of three biological replicates was done for each cell line. CTG assay readouts were plotted as a function of log molar [inhibitor] and a 4-parameter sigmoidal concentration response model was fitted to the data. Mean ± s.d. was plotted for each tested dilution.

### Human cell line proliferation assay

19 PDAC cell lines were tested for sensitivity to RMC-7977 as part of a panel of human cancer cell lines of various histotypes screened at Crown Bioscience. These PDAC cell lines harbored *KRAS^G12D^, KRAS^G12V^, KRAS^G12C^, KRAS^Q61H^, and BRAF^V487_P492delinsA^* mutations. To measure inhibition of cell proliferation, cells were cultured in methylcellulose and treated in triplicates with serial dilutions of RMC-7977 (top concentration of 1 µM) or DMSO dispensed by a Tecan D300e digital dispenser (Tecan Trading AG). Cells were incubated for 120 hours prior to measurement of ATP levels using CellTiter-Glo. Technical triplicates were run for each biological replicate and a total of three biological replicates was done for each cell line. CTG assay readouts were plotted as a function of log molar [inhibitor] and a 4-parameter sigmoidal concentration response model was fitted to the data. Mean ± s.d. was plotted for each tested dilution.

### Western blot analysis

Cells were seeded at 2 × 10^5^ - 4 × 10^6^ cells per well in 6-well plates or 100 mm dishes in growth medium. After overnight incubation, RMC-7977 or DMSO (0.1% v/v) were added and incubated for the indicated time points. Cells were washed twice with ice-cold PBS and lysed with NP-40 lysis buffer (Thermo Fisher Scientific, J60766), MSD Tris Lysis Buffer (MSD, R60TX-2) or a lysis buffer containing 1% Triton X-100, 20 mM Tris-HCl, 150 mM NaCl, and 1 mM EDTA. All lysis buffers were supplemented with protease and phosphatase inhibitors. Lysates were scraped and collected before centrifugation at 21,000 × g for 10 minutes at 4°C. The protein-containing supernatants were quantified by BCA assay (Pierce, 23225) and equal quantities of protein were denatured with LDS and reducing agent at 95°C. Samples were resolved on 12% or 4-12% Bis-Tris polyacrylamide gels, then transferred to a nitrocellulose or PVDF membrane using the iBlot 2.0 system or wet transfer. Membranes were blocked in Intercept TBS buffer (LiCor, 927-60001) or 3% milk before probing with primary antibodies overnight at 4°C. Secondary antibodies were added as appropriate, and the membranes were imaged on a LiCor Odyssey imager. Alternatively, membranes were incubated with HRP-linked secondary antibodies and developed with Clarity or ClarityMax chemiluminescent substrates using a ChemiDoc XRS+ imager (Bio-Rad). (See Supplementary Methods Table 3 for antibody information).

### PDAC organoid preparation and treatment conditions

#### Origins and genetic profiling of patient-derived organoids

Genetic profiling was performed on the patients’ tissue biopsies using whole-genome sequencing or OncoPanel^39,47^. All patients consented to an Institutional Review Board (IRB)-approved protocol at Dana-Farber Cancer Institute permitting access to their clinical and genomic data.

#### Organoid culture

Organoids were cultured at 37°C in 5% CO_2_. Cells were seeded in growth factor reduced Matrigel (Corning; Cat. #356231) domes and incubated with human organoid medium formulated as follows: Advanced DMEM/F12-based-conditioned medium, 1x B27 supplement, 10 mM HEPES, 2 mM GlutaMAX, 10 mM nicotinamide, 1.25 mM N-acetylcysteine, 50 ng/mL mEGF, 100 ng/mL hFGF10, 0.01 μM hGastrin I, 500 nM A83– 01, Noggin 100ng/mL, 1x Wnt-3A conditioned 10% FBS DMEM (50% by volume) and 1X R-spondin Conditioned Basal Medium (10% by volume)^48,49^.

#### Organoid drug treatment and viability assay

Organoids were dissociated using TrypLE Express (Thermo Fisher, Cat. #12604054) and cells were seeded into ultra-low attachment 384-well plates at 1 × 10^3^ cells per well into 20μl of culture media, consisting of 10% Matrigel and 90% human organoid medium. Organoids were treated 24 hours post seeding over a 12-point dose curve with RMC-7977 or with DMSO in a randomized fashion using a Tecan D300e Digital Dispenser. Cell viability was assessed 6 days post-treatment using a Cell-TiterGlo 3D Cell Viability assay (Promega, G9683), according to the manufacturer’s instructions. Fluorescence was read using a FLUOstar Omega microplate reader. Technical triplicates were analyzed for each biological replicate and a total of three biological replicates were done for each cell line. CTG assay readouts were plotted as a function of log molar [inhibitor] and a 4-parameter sigmoidal concentration response model was fitted to the data. Mean ± s.d. was plotted for each dilution.

### *Ex vivo* human PDAC explant preparation and treatment conditions

Explant culture sponges and optimized culture media were prepared as previously described in Hasselluhn and Decker-Farrell et al.^26^. Human tissue samples were obtained from de-identified patients undergoing resection surgeries, primarily pancreaticoduodenectomy (Whipple) or distal pancreatectomy, at New York-Presbyterian/Columbia University Irving Medical Center. Upon receipt of a resected human PDAC fragment (n=4), each tissue sample was cut into 300μm slices using a Compresstome. Any tumor tissue remaining after sectioning was fixed in 4% paraformaldehyde (Santa Cruz Biotechnology, sc-281692) for 2 hours, at 4°C as the Day 0 control. Sectioned slices were next placed between gelatin sponges pre-soaked in 750 µl of media containing either DMSO or varying concentrations of RMC-7977 (10-100 nM). Each well of a 24 well plate contained one explant slice placed between bottom sponge (1 cm^3^) and a top sponge (2-3 mm thick). After 24 hours culture, explants were collected and fixed in 4% PFA for 2 hours. Fixed tissue was then transferred to 70% ethanol and paraffin embedded for long-term storage and further analysis.

### *In vivo* xenograft studies

#### Animal studies

Studies were conducted at the following CROs: GenenDesign (Shanghai, China), Pharmaron (Beijing, China), and Wuxi AppTec (SuZhou, China). All CDX/PDX mouse studies and procedures related to animal handling, care and treatment were conducted in compliance with all applicable regulations and guidelines of the relevant Institutional Animal Care and Use Committee (IACUC). Female BALB/c nude mice and NOD SCID mice 6-8 weeks old from Beijing Vital River/VR Laboratory Animal Co., LTD, Beijing AniKeeper Biotech Co., Ltd., and Shanghai Sino-British SIPPR/BK Laboratory Animal Co., LTD were used for these studies.

#### Generation of xenograft models

In order to generate subcutaneous xenograft tumors each mouse was inoculated at the right flank with tumor cells (2 × 10^6^ - 1 × 10^7^) in 100-200 µl of media/PBS supplemented with Matrigel (1:1). Treatments were started when the average tumor volume reached 150-250 mm^3^ (for tumor growth evaluation) and 400-600 mm^3^ (for single dose pharmacokinetic/pharmacodynamic (PK/PD) study). Tumor diameter was measured in two dimensions using a digital caliper, and the tumor volume in mm^3^ was calculated using the formula: Volume = ((width)2 × length)/2. Mice on studies were weighed and tumors were measured 2 times a week.

The human primary cancer xenograft models were generated using fresh tumor fragments obtained from hospitals with informed consent from the patients in accordance with protocols approved by the Hospital Institutional Ethical Committee (IEC). The tumor fragments were serial passaged in BABL/c nude mice and then cryopreserved for further use. For this study, recovered tumor fragments of about 15-30 mm^3^ in size from each model were implanted into right flanks of BALB/c nude mice. Treatment started when average tumor volume reached 150-250 mm^3^.

To generate orthotopic xenograft tumors, survival surgeries were carried out and 2 × 10^6^ - 5×10^6^ luciferase-expressing tumor cells in 30-50 µL Media/Matrigel mixtures (1:1) were implanted directly into the mouse pancreas. Treatments were started when the tumors produced an average of 50-80 × 10^7^ photon/second as measured by the *in vivo* imaging system (IVIS). All subsequent tumor measures were also conducted by IVIS. For routine monitoring, mice were injected intraperitoneally with 15 mg/mL (at 5 µL/g BW) of D-Luciferin (Perkin Elmer) and imaged, after which Living Image software (Perkin Elmer) was used to compute regions of interest (ROI) and tumor volumes.

#### RMC-7977 treatment

Tumor-bearing animals were randomized and assigned into groups (n=3-10/group). Vehicle or RMC-7977 was administered via oral gavage daily at 10 mg/kg and animals were treated for 21-28 days. Studies were terminated early if tumor burden reached humane endpoint. Body weights were collected twice a week during the study. Means ± s.e.m were plotted in the waterfall plots. For the single-dose pharmacokinetic/ pharmacodynamic (PK/PD) study, mice were randomized and assigned into groups (n=3-6/dose/timepoint). A single dose of RMC-7977 was administered orally at 10 mg/kg, 25 mg/kg and 50mg/kg. Tissues (including tumor, colon and skin) were harvested at indicated time points and either fixed in 10% formalin, embedded in Optimal Cutting Temperature (OCT; Sakura, 4583) solution or snap-frozen in LN2 for further analysis. Whole blood was transferred into K_2_EDTA Microtainer tubes (BD, 365974), incubated for 5 minutes and snap-frozen in LN_2_.

### *In vivo* allograft studies

#### Animal studies

All murine allograft studies and procedures related to animal handling, care and treatment were conducted in compliance with all applicable regulations and guidelines of the Institutional Animal Care and Use Committee (IACUC). Female C57BL/6J (strain 000664) mice aged 6-8 weeks from the Jackson Laboratory (Bar Harbor, ME USA) were used for these studies.

#### Generation of allograft models

In order to generate subcutaneous (SC) allograft tumors, each mouse was inoculated in the right flank with 3 × 10^5^ of KPCY 6499c4 tumor cells in 0.1 ml of Matrigel:PBS (1:1). Treatments were started when the average tumor size reached 140 mm^3^. Tumor size was measured at two dimensions using a digital caliper, and the tumor volume in mm^3^ was calculated using the formula: Volume = ((width)2 × length)/2. Mice on studies were weighed and tumors were measured 2 times a week.

To generate orthotopic allograft tumors, 5 × 10^4^ KPCY 6499c4 tumor cells in 20 µL PBS/Matrigel mixtures (1:1) were implanted directly into the mouse pancreas through a laparoscopic incision. Treatments were started when the average tumor size reached ∼50 mm^3^. Body weights were measured and tumor growth was monitored by ultrasound twice weekly.

#### RMC-7977 treatment

Tumor-bearing animals were randomized, assigned into groups (n=9-10/group), and treated daily via oral gavage with Vehicle or RMC-7977 (10 mg/kg, q.d.). For SC KPCY study, survival endpoint was defined as: tumor volume reaching 2000 mm^3^ or mice showing any clinical signs, including severe ulceration. For orthotopic KPCY study, survival endpoint was defined as (1) mice showing any clinical signs including hunching or fluid in the abdomen, or (2) tumor dimensions exceeding the imaging frame of the ultrasound. Body weights were measured twice a week during the study. Tissue was harvested either at 4 hours or 24 hours after last dose and preserved as previously described (See Xenograft studies section).

### *In vivo* GEMM studies

#### Animal Breeding

All animal research experiments were approved by the Columbia University Irving Medical Center (CUIMC) Institutional Animal Care and Use Committee (IACUC). Mouse colonies were bred and maintained with standard mouse chow and water, ad libitum, under a standard 12hr light/12hr dark cycle. KPC (*Kras^LSL.G12D/+^*; *p53^LSL.R172H/+^*; *Pdx1-Cre*), KC (*Kras^LSL.G12D/+^*; *Pdx1-Cre*), PC (*p53^LSL.R172H/+^*; *Pdx1-Cre*) as well as KP^f/f^C (*Kras^LSL.G12D/+^*; *p53^flox/flox^*; *Pdx1-Cre*), KP^f/f^ (*Kras^LSL.G12D/+^*; *p53^flox/flox^*) and P^f/f^C (*p53^flox/flox^*; *Pdx1-Cre*) mice were generated in the Olive Laboratory at Columbia University, by crossing the described alleles. Mouse genotypes were determined using real time PCR with specific probes designed for each gene (Transnetyx; Cordova, TN). *Kras*^LSL-G12D/+^; *p53*^LSL-R172H/+^, *Pdx1-Cre*, *Rosa26*^YFP/YFP^ (KPCY) were bred and maintained in pathogen-free facilities at the University of Pennsylvania.

#### Pharmacokinetic/pharmacodynamic study in KP^F/F^C

Tumor formation in KP^F/F^C *(Kras^LSL.G12D/+^*; *p53^flox/flox^*; *Pdx1-Cre*) mice was monitored by bi-weekly palpations until the detection of a mass, which was then confirmed by ultrasound. Tumor-bearing mice were randomized and assigned into groups (n=3/dose/timepoint). Single dose of RMC-7977 was administered orally at 10 mg/kg, 25mg/kg and 50mg/kg. The same vehicle formulation was used for the control group. Whole blood and tissue (tumors and colons) were harvested at indicated time points preserved as previously described (See Xenograft studies section).

#### Pharmacodynamic study in KPC mice

Tumor formation in KPC mice was monitored by bi-weekly palpation. Upon detection of a 4–7 mm diameter tumor by ultrasound, KPC mice were randomized and treated with Vehicle (n=6) or RMC-7977 (50 mg/kg; n=8). Treatments were performed every other day via oral gavage for 1 week. Animal health status and weight were checked daily and ultrasounds (Vevo 3100) were performed every third day to monitor tumor growth. Following two consecutive ultrasounds, RMC-7977-treated animals were sacrificed either 4 (n=4) or 24 hours (n=4) after last dose and Vehicle-treated animals were sacrificed between 4-24 hours post last dose. Tissue was harvested and preserved as previously described (See Xenograft studies section).

#### Pharmacodynamic study in KPCY mice

KPCY mice were enrolled upon detection of a 15-100 mm^3^ tumor measured via ultrasound. Animals were randomized into groups and treated with Vehicle (n=6) or RMC-7977 (25 mg/kg; n=8). Treatments were performed every day via oral gavage for 15 days and ultrasounds were performed on day 8 and 15. Animals were sacrificed after last dose and tissue was collected and preserved as previously described (See Xenograft studies section).

#### Survival study in KPC mice

For survival study, KPC mice with 4–7 mm diameter tumors were enrolled and treated every other day with Vehicle (n=8) or RMC-7977 (50 mg/kg; n=8). Animal health status and weight were checked daily and ultrasounds were performed every third day to monitor tumor growth. Survival endpoint was defined as mice showing any clinical signs including hypothermia, hemorrhagic ascites or GI obstruction. Mice that reached endpoint criteria were sacrificed either at 4 or 24 hours after last dose and tissue was collected for further analysis.

### *In vivo* pharmacodynamic analysis by *DUSP6* qPCR

RNA was extracted from at least 20 mg of tissue using an RNeasy Mini Kit (Qiagen, 74104) and a High Throughput Tissue grinder following the manufacturer’s protocol. Reverse transcription was carried out using High-Capacity cDNA Reverse Transcription Kit (ABI, 4368814) according to the manufacturer’s protocol. The cDNA product was used for qPCR analysis using TaqMan Gene Expression Master Mix (ABI, 4369016). TaqMan primer probes specific to *DUSP6* (human - Hs00737962_m1, murine - Mm00518185_m1, FAM-MGB) and *18S* (human - Hs99999901_s1, murine - Mm03928990_g1, FAM-MGB, used as an internal control gene) were used to detect the levels from each sample in duplicates using a 10 µL final reaction volume in a 384-well clear optical reaction plate. For qPCR, Ct value of *DUSP6* and *18S* were obtained for analysis. *DUSP6* Ct value was normalized to *18S*, and then the mean relative mRNA expression levels of each group were normalized to the vehicle control group. Values were plotted as relative change in mRNA expression compared to Vehicle. Means ± s.d. were shown.

### Mouse blood and tissue sample bioanalysis

Whole blood, tumor, colon and skin tissue concentrations of RMC-7977 were determined using liquid chromatography-tandem mass spectrometry (LC-MS/MS) methods. Tissue samples were homogenized with a 5× or 10× volume of homogenization buffer (methanol/15 mM PBS (1:2; v:v) or 15 mM PBS with 10% methanol). An aliquot of whole blood or homogenized tissue (10 or 20 µL) was transferred to 96-well plates (or tubes) and quenched with a 20× volume of acetonitrile/methanol (1:1; v/v) with 0.1% formic acid containing a cocktail of internal standards (IS). After thorough mixing and centrifugation, the supernatant was directly analyzed on a Sciex 6500+ triple quadrupole mass spectrometer equipped with an ACQUITY or Shimadzu UPLC system. An ACQUITY UPLC BEH C18 or C4 1.7 μm (2.1 × 50 mm) column was used with gradient elution for compound separation. RMC-7977 and IS (verapamil or terfenadine) were detected by positive electrospray ionization using multiple reaction monitoring (RMC-7977: m/z 865.4/706.4 or m/z 865.3/833.5; verapamil: m/z 455.2/164.9; terfenadine: m/z 472.3/436.4). The lower limit of quantification was 0.5 ng/mL or 2.0 ng/mL for blood, tumor, and other tissue. BA analysis on blood and tissue samples from xenograft models was run at Wuxi AppTec. BA analysis on blood and tissue samples from allograft models and GEMM was run at Revolution Medicines.

### Immunohistochemistry (IHC)

All tissues were fixed for up to 24 hours using 10% neutral buffered formalin and then moved to 70% ethanol for long term storage. All stainings were performed on 4-µm tissue sections. Sections were deparaffinized using a Leica XL ST5010 autostainer, after which slides were subjected to heat-activated epitope retrieval. To block endogenous peroxidases, 20 min incubation in 3% H_2_O_2_ (Fisher Scientific) was performed. Slides were further blocked in serum for 1 hour, and primary antibodies were added for overnight incubation at 4°C. The next day, slides were washed and incubated with ImmPRESS HRP Horse Anti-Rabbit IgG Polymer Detection Kit (Vector Laboratories, MP-7401) for 30 min. Following incubation, ImmPACT DAB peroxidase (Vector Laboratories, SK-4100) was used to develop the stain and hematoxylin was used as nuclear counterstain. Stained slides were imaged at 40x magnification. Quantitative analyses of IHC images were performed using ImageJ.

To stain tissues collected from the Capan-1 xenograft model, a similar protocol was used with the following changes. Sections were stained using a Leica BOND automated staining system and primary antibodies were detected with the Leica BOND Polymer detection kit (3-P-PV6119). Stained slides were scanned and digitized with a 3DHistotech Pannoramic whole slide scanner at 20x magnification. Image analysis was performed using HALO software from Indica Labs.

To stain tissues collected from the KPCY allograft model, a Biocare IntelliPATH automation system was used, and primary antibodies were detected with the MACH4-HRP-polymer Detection System (Biocare, MRH534). Stained slides were scanned and digitized with a TissueScope LE (Huron Digital Pathology) whole slide scanner at 20x magnification. Image analysis was performed using HALO software from Indica Labs. (See Supplementary Methods Table 3 for antibody information).

## Acknowledgements

This study was funded in part by Revolution Medicines. The authors thank Crown Bioscience and the PRISM lab for implementation of in vitro assays and GenenDesign, Pharmaron, Wuxi AppTec for animal study support. The authors thank Anirban Miatra (M.D. Anderson Cancer Center) for PDAC cell lines. C.J.D. received support from the National Institutes of Health (NIH; R01CA42978, R01CA175747, R01CA223775, P50CA196510, U01CA199235 and P01CA203657 and R35CA232113), the Pancreatic Cancer Action Network/AACR (15-90-25-DER), the Department of Defense (W81XWH-15-1-0611), and the Lustgarten Foundation (388222). K.D.M. received support from the NIH (T32CA009156) and American Cancer Society (PF-22-066-01-TBE). A.J.A. received support from the NIH (R01CA276268) and the Lustgarten Foundation, Break Through Cancer, and Hale Center for Pancreatic Cancer Research. B.Z.S. and R.H.V. received support from the NIH (R01CA229803). S.B.K. received support from CRI Irvington Postdoctoral Fellowship. GAM received support from the NIH (1R01CA266558, 1U01CA274312). A.C.G. received support from the Charles H. Revson Senior Fellowship in Biomedical Science (Grant No. 22-22). K.P.O. received support from the NIH (1R01CA266558, 2R01CA215607, 1U01CA274312), a Translational Clinical Program Grant from the Lustgarten Foundation (17-15), and the Rockhammer Charitable Fund. This work was supported by the HICCC Cancer Center Support Grant (P30CA013696) and DLDRC (P30 DK132710) and included use of the following shared resources: Genomics and High Throughput Screening, Oncology Precision Therapeutics and Imaging, and Molecular Pathology. We are grateful to members of the CONTrasT Consortium for their comments and suggestions.

## Author contributions

Authors contributed as followed: Conceptualization – U.N.W., J.J., A.J.A., B.Z.S., R.H.V., C.J.D., Z.W., M.S., K.P.O.; Resources – B.B., J.A.C., M.D.K., G.A.M. D.W., J.A.M.S., E.Q., A.J.A., B.Z.S., R.H.V., C.J.D., M.S., K.P.O.; Data curation – U.N.W., J.J., Y.W., M.M., B.L., J.D., M.O., K.D.M., M.H., S.C., C.H., Y.G., Y.C.Y., R.Z.; Software – Y.G., R.Z., A.C.; Formal analysis – U.N.W., J.J., Y.W., M.M., B.L., S.C., R.Z., C.H., Y.G., Z.W.; Supervision – J.J., D.W., J.A.M.S., E.Q., A.J.A., B.Z.S., R.H.V., C.J.D., M.S., K.P.O.; Validation – U.N.W., J.J., M.M., M.O., J.D., K.D.M., M.S., K.P.O.; Investigation – U.N.W., J.J., T.D., M.C.H., A.C.G., C.P., S.S.A., M.O., K.D.M., J.D., S.B.K., R.S., W.V., A.A.A., P.S.H., C.H., M.H., M.S.; K.P.O.; Visualization – U.N.W., J.J., Y.W., M.M., B.L., S.C., Y.G., J.D., M.O., K.D.M., Z.W., M.S., K.P.O.; Methodology – Y.W., M.M., B.L., S.C., Y.G., A.R.D.F., S.B.K.; Project administration – J.J., Y.W., M.M., B.L., L.J., X.W., S.C., Y.G., Y.C.Y., R.Z., H.C.,C.H., C.C., A.J.A., B.Z.S., R.H.V., C.J.D., K.P.O.; Writing of the original draft – U.N.W., J.J., M.A.B., M.S., K.P.O.; Writing-review and editing – U.N.W., J.J., M.O., J.D., K.D.M., A.A.A., P.S.H., C.H., A.C.G., T.D., A.R.D.F., M.C.H., M.A.B., M.H., D.W., J.A.M.S., A.J.A., B.Z.S., R.H.V., C.J.D., M.S., K.P.O.. Contributions were structured using the CREDIT definitions from https://credit.niso.org/.

## Additional information

Raw data for Main and Extended figures as well as Extended Data Tables were submitted with this manuscript.

## Supplementary information

Supplementary Methods Tables were submitted with this manuscript.

---

**Extended Data Figure 1.**
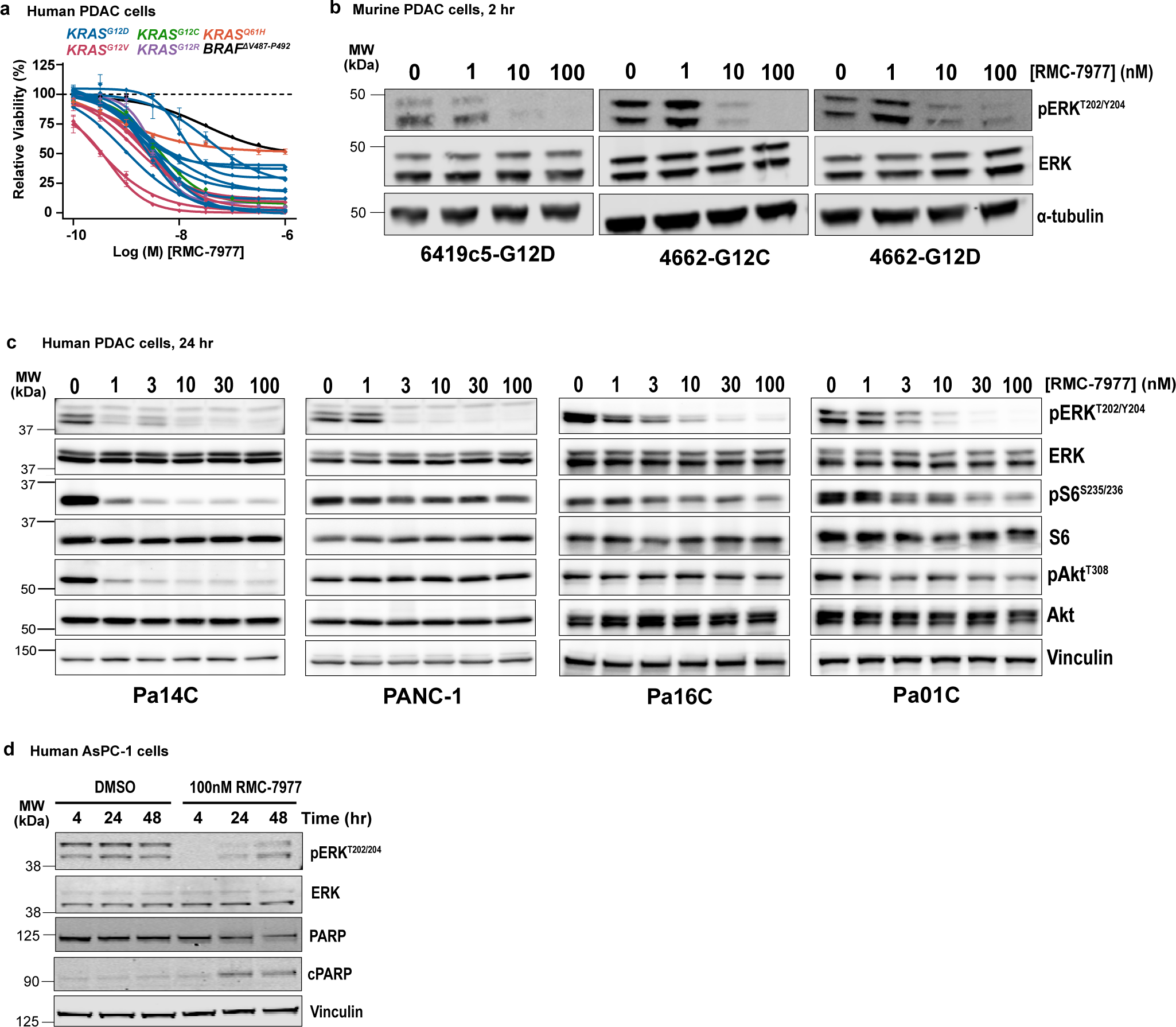
**(a)** Viability levels of human PDAC cell lines with KRAS^G12X^ (AsPC-1, HPAC, Panc 05.04, PANC-1, Panc 10.05, PL45, SU.86.86, SW-1990, KP-4, HPAF-II, Capan-1, Capan-2, CFPAC-1, Panc 03.27, HuP-T4, MIA PaCa-2 and PSN-1), KRAS^Q61H^ (Hs 766T), and BRAF^ΔV487^–^P492^ (BxPC-3) mutations treated with indicated concentrations of RMC-7977 for 4 hours. Data points represent the mean of technical replicates normalized to DMSO control. Error bars indicate s.d. KRAS mutations are indicated by curve colors. **(b)** Representative Western Blot images for three murine PDAC cell lines treated with DMSO or range of RMC-7977 concentrations (1-100 nM) for 2 hours. Protein levels of phospho-ERK_T202/204_ and total ERK were analyzed. Alpha-(α)-tubulin was used as loading control. **(c)** Representative Western Blot images for human PDAC cell lines treated with DMSO or range of RMC-7977 concentrations (1-100 nM) for 24 hours. Protein levels of phospho-ERK^T202/204^, total ERK, phospho-pS6^S235/236^ total S6, phospho-Akt^S473^ and total Akt were analyzed. Vinculin was used as loading control. **(d)** Representative Western Blot images for AsPC-1 cell line treated with DMSO or RMC-7977 (100 nM) for indicated timepoints. Protein levels of phospho-ERK^T202/204^, total ERK, total PARP and cleaved PARP were analyzed. Vinculin was used as loading control.

**Extended Data Figure 2.**
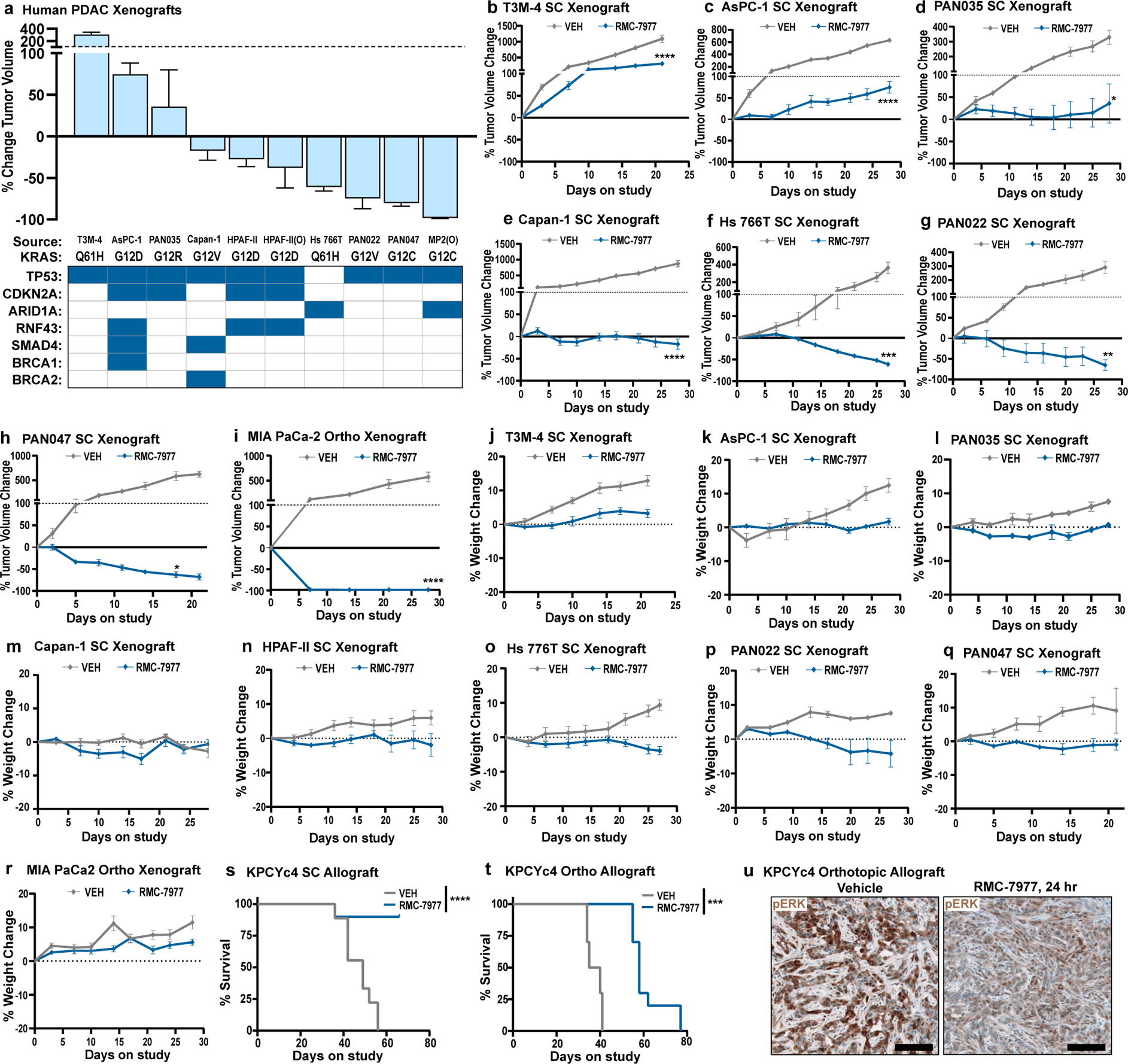
**(a-k)** Human PDAC xenograft models from Fig. 2. **(a)** Waterfall plot showing tumor volume change from baseline in RMC-7977 treated tumors. Error bars indicate ± s.e.m. Table shows selected genotypes for the xenograft panel with the row above indicating the KRAS mutation. Present co-mutations in each model shown as blue squares in the table. **(b-i)** Tumor growth curves for indicated xenograft models from (a), shown as percent tumor volume change from baseline over the course of treatment. Vehicle and RMC-7977 groups were compared by 2-way repeated measures ANOVA on the last measurement day of the vehicle group (*, p<0.05; **, p<0.01; ***, p<0.001; ****, p<0.0001). Error bars indicate ± s.e.m. **(j-r)** Tolerability of RMC-7977 as assessed by percent animal body weight change from baseline over the course of treatment, for indicated xenograft models from (a). Error bars indicate ± s.e.m. **(s,t)** Kaplan-Meier survival analysis comparing RMC-7977 and Vehicle treatment arms in **(s)** subcutaneous and **(t)** orthotopic KPCYc4 allograft models (***, p<0.001; ****, p<0.0001). **(u)** Representative IHC images of Vehicle and RMC-7977-treated KPCYc4 tumors collected at 24 hours post last dose and stained for phospho-ERK^T202/204^. Scale bars = 100 μm.

**Extended Data Figure 3.**
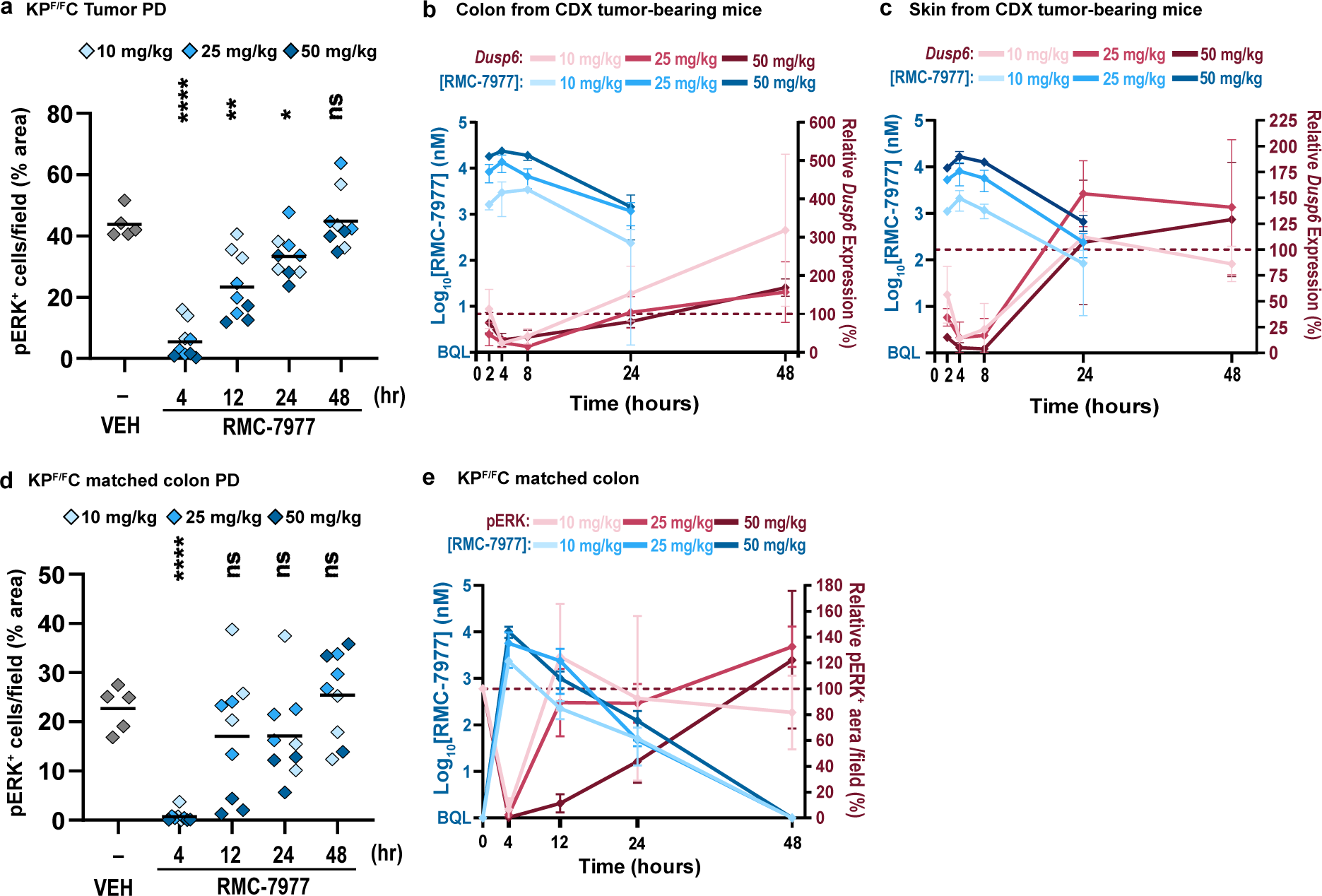
**(a,d)** IHC analysis was ran on tissues collected from KP^F/F^C mice in Fig 3. **(a)** Quantification of phospho-ERK^T202/204^ IHC staining of tumors isolated from Vehicle and RMC-7977 treated KP^F/F^C mice. **(b,c)** PK/PD relationship in the colons and skin isolated from CDX tumor-bearing mice. Pharmacokinetic response shown as RMC-7977 concentration in **(b)** colon and **(c)** skin (solid blue lines) over time. Pharmacodynamic response shown as relative change in *Dusp6* expression in colon or skin (red solid lines) over time. Shades of blue or red represent three tested doses. Error bars indicate s.d. **(d)** Quantification of phospho-ERK^T202/204^ IHC staining of colons isolated from Vehicle and RMC-7977-treated KP^F/F^C mice. **(e)** PK/PD relationship of RMC-7977 in the colons isolated from KP^F/F^C mice. Pharmacokinetic response shown as RMC-7977 concentration in colon (solid blue lines) over time. Pharmacodynamic response shown as relative change in pERK positive IHC staining in colon (red solid lines) over time. Shades of blue or red represent three tested doses. Error bars indicate s.d. **(a,d)** Analysis of IHC images based on 10-15 fields of view and plotted as average per tissue section. Shades of blue represent three tested doses. Results were compared by Student’s unpaired t-test (*, p<0.05; **, p<0.01; ****, p<0.0001).

**Extended Data Figure 4.**
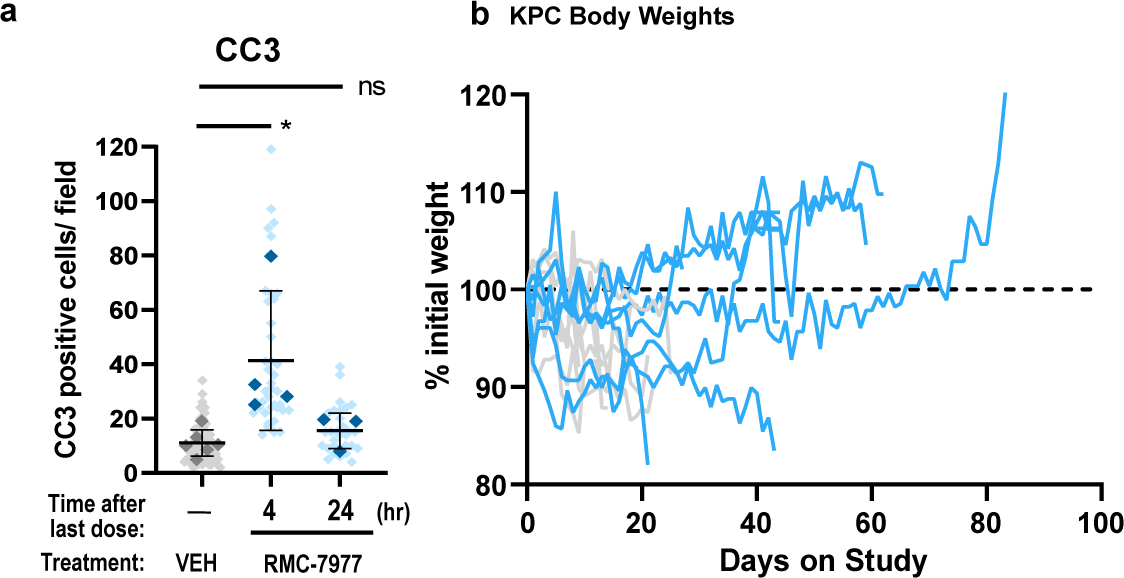
**(a)** IHC analysis of KPC tumors from Fig. 4. Tumors stained for CC3. Quantification of IHC images was compared between Vehicle, RMC-7977 (1 week + 4 hour) and RMC-7977 (1 week + 24 hour). Quantification was based on 10-15 fields of view (light shade), averaged per tumor (dark shade) and means were compared by Student’s unpaired t-test (*, p<0.05). Error bars indicate s.d. **(b)** Tolerability of RMC-7977 in KPC mice as assessed by percent animal body weight change from baseline over the course of treatment.

